# A fast microfluidic device to measure the deformability of red blood cells

**DOI:** 10.1101/644161

**Authors:** Ninad Mehendale, Dhrubaditya Mitra, Debjani Paul

## Abstract

We report a microfluidic device to determine the shear elastic modulus and the Young’s modulus of red blood cells (RBCs). Our device consists of a single channel opening into a funnel, with a semi-circular obstacle placed at the mouth of the funnel. As a RBC passes the obstacle, it deflects from its original path. Using populations of artificially-stiffened RBCs, we show that the stiffer RBCs deflect more compared to the normal RBCs. We use calibration curves obtained from numerical simulations to map a trajectory of each RBC to its elastic constants. Our estimates of the shear elastic modulus and the Young’s modulus of normal RBCs are within the same range of values reported in the literature using AFM, optical tweezers and micropipette measurements. We also estimate indirectly the elongation index of normal and artificially hardened RBCs from their tracks, without any direct observation of their shapes. Finally, we sort a mixed population of RBCs based on their deformability alone. Our device could potentially be further miniaturized to sort and obtain the elastic constants of nanoscale objects, whose shape change is difficult to monitor by optical microscopy.

## 1. Introduction

Red blood cells (RBCs) are more deformable (i.e., shear elastic modulus **∼** 2 - 10 μN/m)^1,2^compared to the most other cells in the body, which allows them to transport oxygen through the microcirculation. Any change in the deformability of RBCs can have major physiological effects, e.g., if the RBCs become stiffer, they can occlude the blood flow in capillaries and eventually lead to excessive RBC destruction in the spleen. The stiffness of the RBCs changes under certain pathophysiological conditions, such as, malaria^3^, sickle cell disease^4^, diabetes^5^, peripheral vascular disease^6^, etc. This, in turn, gives rise to the possibility that such conditions can be detected by measuring the elastic properties of RBCs. As RBCs are heterogeneous biological objects, we expect their physical parameters to be distributed over a range of values. For example, the variation in the size of RBCs for a single individual is given by the RBC distribution width. Any change in the distribution width could signify a pathophysiological condition^7^. Clearly, we first need to determine the distribution of elastic moduli (e.g. Young’s modulus, shear modulus, etc.) of RBCs in healthy individuals before we can proceed to understand why and how they change in disease. This kind of information can only be obtained from single-cell measurements performed on a large population of RBCs.

There are a number of conventional techniques by which the deformability of the RBC population can be measured at the level of single cells: e.g., micropipette aspirations^8^, atomic force microscopy (AFM)^9,10^, membrane fluctuations^3,11^, optical tweezers^12^, optical stretcher^13^, etc. Most of these techniques require expensive specialized equipment, have a low throughput and the data acquired by these methods takes a long time to analyze. These challenges led to the exploration of microfluidic techniques. Most microfluidic techniques do not directly measure the elastic constants, but rely on accurate monitoring of the shape change of RBCs in real time as they pass through a constriction. In most cases the change in shape is parametrized with one or more parameters which are used as surrogates of the elastic coefficients. In one case ^14^, the change in shape is compared to the numerical calculation of a model to derive an elastic constant. This approach, while fast, still requires expensive high-speed cameras (2000 - 4000 fps) to track shape changes of cells in real time. Another class of microfluidic techniques measures the pressures required to pass the RBCs through constrictions narrower than their size. The cortical tension of the RBCs is determined from these measurements^15^.

Here we report a microfluidic device that measures the shear elastic modulus (*Gs*), the Young’s modulus (*Y*) and the elongation index (*E. I.*) of a large number of single RBCs within a few minutes by deflecting them from their paths using an obstacle placed in a laminar microfluidic flow. Instead of using a high-speed (3000 **–** 4000 fps) camera to track the shape change of the RBC in real time, we track its path using a regular 25 fps CCD camera. We then combine our experimental observations of RBC tracks with COMSOL simulation to obtain the shear elastic modulus (*G*_*s*_) the Young’s modulus (*Y*) and the Elongation Index (*EI*) of each individual RBC. By performing this experiment with a large population of RBCs, we can obtain the distribution of the elastic moduli of the entire RBC population. This technique of measuring the cell elasticity is both cheaper and simpler compared to the reported microfluidic methods as we do not need to fabricate devices with very small feature sizes (i.e. comparable to the lateral dimension of RBCs, which is ∼ 2 *μ*m or less) or use high-speed (i.e. several thousands of fps) cameras. We can also accurately determine the distribution of the elastic constants of a mixed population of RBCs consisting of deformable and stiff cells. Furthermore, the observation that normal and stiff RBCs exit the device at different angles suggests the potential of our device for sorting of cells based on their mechanical properties. The device can be, in principle, be scaled down by a factor of hundred to measure and detect the deformability of extracellular vesicles, which are so small that they cannot be resolved under the optical microscope.

## 2. Results and discussions

### 2.1 gh1 RBCs of different deformability follow different trajectories after passing the obstacle

Based on direct numerical simulations, Zhu et. al. ^16^ proposed a microfluidic device, which is somewhat similar to the Rutherford scattering experiment, to separate capsules by their deformability. The main feature of this microfluidic device is a single semi-cylindrical (semi-circular in 2D view) obstacle placed at the mouth of a funnel (fig. 1A). In their numerical simulations, capsules with different stiffnesses are made to flow towards the obstacle. These capsules deform while going past the obstacle **–** soft capsules deform more than the stiff ones **–** and consequently follow different trajectories in the funnel depending on their deformabilities, eventually exiting the device at different locations. Stiffer capsules deviate more from their initial track compared to the softer capsules. The purpose of the funnel after the obstacle is to magnify the small differences in the trajectories of the capsules of different stiffnesses as they go past the obstacle. Vesperini, et. al. ^17^ experimentally demonstrated the separation of highly polydisperse vesicles based on size and deformability using the same device design. They showed that at low flow rates the capsules are separated based on size (similar to pinch flow fractionation), while at high flow rates they are separated by stiffness. They also suggested a critical capillary number (between 0.1 and 0.2) to distinguish between these two sorting regimes.

**Figure 1.**
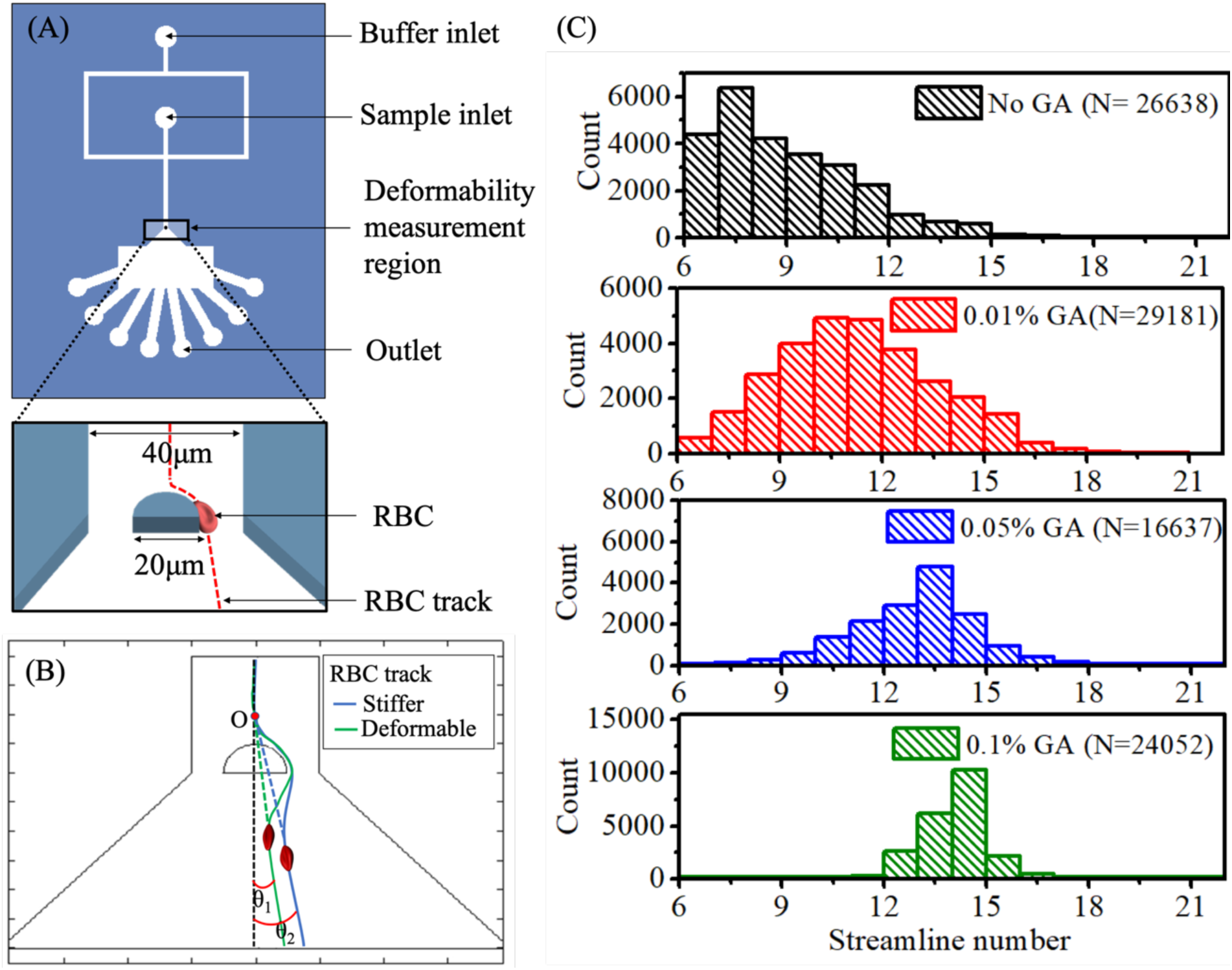
Schematic diagram of the microfluidic device to deflect normal and stiff RBCs from their paths. (A) The device has a 40 μm-wide straight channel that opens into a funnel. A semi-circular obstacle of 20 μm diameter is placed at the mouth of the funnel. A flow-focused and very dilute stream of RBCs moves along the mid-plane of the straight channel towards the obstacle, allowing only one RBC to approach the obstacle at a time. (B) Softer RBCs are deflected by a smaller angle (θ_1_) while stiffer RBCs are deflected by a larger angle (θ_2_) in this device geometry. (C) Histograms showing the tracks of different RBC populations. Each panel denotes a RBC population treated with a specific concentration of a chemical stiffening agent glutaraldehyde (GA).N indicates the number of RBC tracks analysed to obtain a particular distribution. The data is not normalized. Untreated RBCs follow the streamlines with lower numbers, while the progressively stiffer RBCs are found along the streamlines with higher numbers.

We use the same device design (see fig. 1A and 1B), but go further, by sending red blood cells (RBCs) through this device. We first flow a solution of normal healthy RBCs along a short and straight channel segment towards the obstacle. After crossing the obstacle, different RBCs follow different trajectories and exit the device at different angles. Simulations of Zhu and others^16^ suggest that after an RBC passes the obstacle, its centre of mass follows a streamline of the background flow. Hence, to label the tracks of the RBCs, we solve, numerically, the background flow (i.e. the flow without any RBC) with a grid resolution that is consistent with the resolution of our camera. Then we number the streamlines obtained from our simulation from 0 at the center of the funnel to 50 on each side. We track the position of each RBC as it moves along the funnel after passing the obstacle to identify which streamline it follows. We count the number of the RBCs moving along each streamline and plot the data as a histogram in fig. 1C (top panel). Next, we use several batches of RBCs **–** we treat each batch with a specific concentration (0.01%, 0.05% and 0.1% respectively) of a chemical stiffening agent glutaraldehyde (GA) to generate artificially stiffened populations of RBCs with different mean stiffness values **–** and repeat the same experiment. GA is widely used to attach RBCs to different surfaces and also to reduce their deformability^18^. The successive panels in fig. 1C show the positions of the RBCs along different streamlines when different concentrations of GA are used. In the next section we show how we use this data to obtain the deformability distribution of the RBC populations, both the normal and the progressively stiffened ones.

### 2.2 Measurement of the deformability of RBCs

At this point, recall what we understand by the ‘deformability’ of a single RBC. As the simplest example, consider a three-dimensional, homogeneous and isotropic material. For small deformations, a complete description of the elastic properties of this material is given by two scalars (for example, the two Lamé coefficients). Since a RBC is a complex biological object, it is not obvious how many independent scalars are necessary to describe completely its elastic properties. Measurements of the elastic properties of RBC have a long history of using different methods, e.g., AFM, micropipette aspiration, optical tweezers, membrane fluctuations, etc. Each technique may end up measuring a different aspect of the deformability of RBCs. For example, AFM measurements yield elastic constants at length scales of the order of the AFM tip (∼ nm), while micropipette measurements can probe the elastic properties at a larger length scale (∼*μ*m). Hence, it is not possible to directly compare the results of these techniques with each other unless different methods are cross-calibrated against each other using the same RBC^19^. Finally, the results of a particular class of measurements are interpreted using several different models of the RBC, which makes the measured constants model-dependent.

One class of measurements attributes the results completely to the elasticity of the RBC membrane, while ignoring the inner constituents of the cell. These techniques measure different elastic constants of the membrane: (a) shear modulus (measured by micropipette aspiration^8^, optical tweezers^12^, magnetic twisting cytometry^20^, and membrane fluctuations^3^), (b) bending modulus (given by micropipette aspiration^21^ and membrane fluctuations^3^), (c) area extension modulus measured using micropipette aspiration^22^, and (d) the aspect ratio^23^ of the RBC, which is not an elastic modulus but can be used as a somewhat indirect measure of deformability. Another class of measurements attributes their results to the entire RBC as a solid object. They interpret their results in terms of the Young’s modulus measured using an AFM^9,24^. Finally, there is a class of experiments^25,26^ that report the transit times or the velocities of soft or stiff RBCs through a constriction as surrogate measures of their deformability. Since these measurements do not yield an actual elastic modulus, we will not discuss those in this paper.

We interpret the data obtained from our experiments in two different ways to obtain the distributions of the shear modulus and the Young’s modulus of RBCs. We also obtain the elongation index of the cells without any direct observation of their shapes. In each case, we need a calibration curve that uniquely maps the trajectory of a RBC to a measure of its deformability. By limiting the height of our chamber to 5 *μ*m, we prevent the RBCs from bending -- hence we cannot measure their bending modulus. We assume that the area of the RBC during deformation remains constant in our experiment. Hence, we also cannot measure the area extension modulus.

To construct the calibration curve, it would have been ideal if we could design capsules with the same shape as an RBC, but with a membrane whose elastic constant is known by other measurements. As this is impossible at this point, we must resort to numerical simulations. In principle, the trajectory of an RBC through our device can be found by solving the flow equations outside the cell -- the Navier-Stokes equations (not Stokes as our Reynolds number is about 10) -- the partial differential equations of elasticity on the cell-membrane -- whose position itself is determined by the solution -- and equations of flow inside the cell. This would require a detailed model of both the cell-membrane and the inside of the cell. Together, this is a very formidable problem^27,28^; more so because the models of RBC membrane itself are not well known in every detail. We have navigated this swamp by a patchwork of several simplifying assumptions. Note first that the device is constructed such that the RBCs lie flat -- they do not have the freedom to turn sideways -- while they go through the device. Because of this strong confinement we have ignored the dimensions perpendicular to the diagram shown in fig 1A and B. This is our first assumption. Second, we have assumed the RBCs to be circles when seen from the top -- the experimental setup makes sure that they always lie in the plane of fig 1A -- with a shear-modulus *G*_*s*_ The area-modulus of the RBC is several orders of magnitude higher than its shear modulus ^29^, hence we have assumed that its area is unchanged in the device, and it is incompressible.

We visualize the motion of the RBC through the device the following way: as the RBC approaches the obstacle it slows down and gets more and more deformed; when it is just about to go through either one of the gaps it is temporarily stationary and in its most deformed state; after which it flows through the gap. We assume that at this last stage of its movement, it essentially follows a streamline of the underlying flow (calculated by solving the Navier--Stokes equations with the geometry of the device, but not the RBC, in COMSOL). The position of this streamline is determined by the RBC’s most deformed shape -- in particular it is the streamline that passes through its center-of-mass at the moment when it is most deformed. From optical measurements, we can reconstruct the track of an RBC; this track has a finite width Δ*l*. This is the uncertainty in our knowledge of the track of the RBC. We use COMSOL to solve the underlying flow (see fig. 2A). Then we calculate the streamlines of the flow that pass through the outlet of our device at every Δ*l* starting from the center of the outlet. These streamlines are numbered from 0 (center of the outlet) to 50 (increasing on both sides). Given one such streamline, we can now identify the position of the center-of-mass of the most deformed shape of the RBC that passes through this streamline (see the section on video processing in Materials and Methods for details). As we consider the problem in two dimensions, the most deformed shape of the RBC is a two-dimensional area, two sides of which, we assume, have the same curvature as the obstacle (fig. 2B). We can estimate this shape by equating its area to the area of the undeformed RBC. Since the flow is incompressible, there must always be a small layer of fluid between the RBC and the obstacle. We assume that this layer has the thickness of four grid points in the highest resolution of our COMSOL simulations. At this point, we are able to map every trajectory to the shape and the position of the RBC at its most deformed state. Note that at this point the RBC is also momentarily stationary, hence the boundary condition of the flow velocity at the surface of the RBC is no-slip. Now we put an additional rigid object in our second COMSOL simulation at the estimated position of the most deformed RBC and with a shape that is the estimated shape of the RBC (see fig 2C). From the solution of the flow problem we can calculate the pressure around the RBC (*P*_1_ and *P*_2_) in its most deformed shape. These pressure values are then used to calculate both the shear modulus (*G*_*s*_) and the Young’s modulus (*Y*) as explained in the following sections.

**Figure 2.**
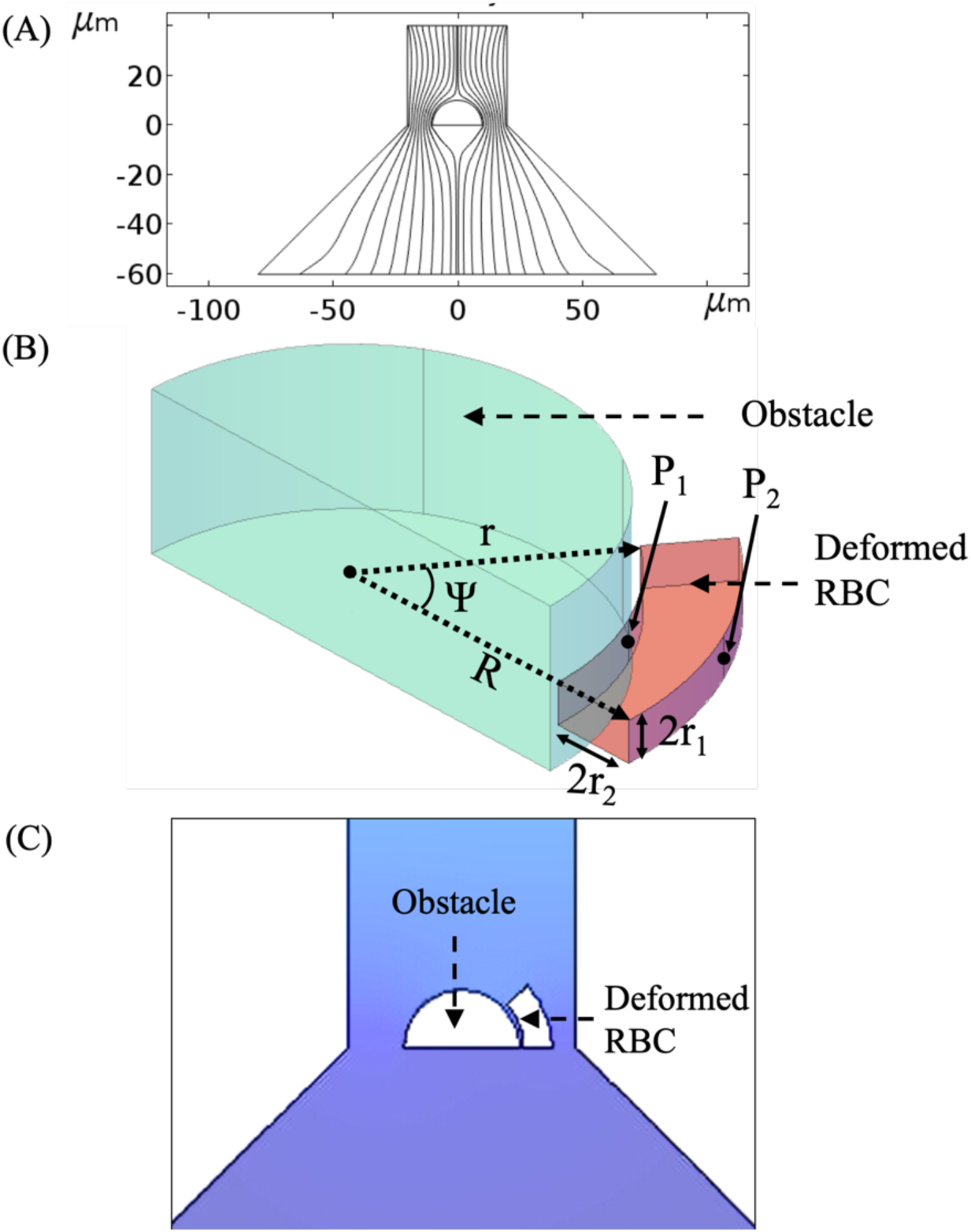
Simulations to obtain the calibration curves mapping different elastic moduli uniquely to the streamline numbers. (A) Streamlines obtained by solving the equations for the background flow in absence of RBCs in the device. These are numbered from 0 to 50, starting from the central streamline. The track of each RBC is labelled by the number of the streamline its centre of mass lies upon. For clarity, every 8th streamline is shown in this image. (B) Schematic diagram showing the obstacle and a shape representing the deformed RBC, where, R and r are its outer and inner radii of curvature. The deformed width is given by 2r_2_ while the height of the RBC is given by 2r_1_The dimensions and the pressure values are used to calculate the Laplace pressure across the RBC membrane. (C) Setup of the second flow simulation in presence of the deformed RBC to obtain P_1_ and P_2_The position of the RBC on the streamline is obtained by combining the data from our experiment and the simulation of the background flow.

#### 2.2.1 Measurement of the shear modulus

To calculate the shear modulus, we assume that the pressure difference (fig. 2B) across the membrane of the RBC is balanced by the Laplace pressure of the membrane. This gives:

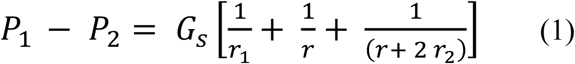

Where *P*_1_ and *P*_2_ are the pressures on the two sides of the bent RBC calculated from our simulations. The values of *P*_1_ and *P*_2_ are obtained from the simulation setup shown in fig. 2C. Equation 1 allows us to calculate the calibration curve (fig. 3A) which maps each streamline to a single value of *G*_*s*_ From our experiment, we can thus generate a plot of the probability density function (PDF) of *G*_*s*_, as shown in fig. 3B. From this plot, we find that the “_#_ for the normal RBCs (black curve) peaks at **∼** 4.8 *μ*N/m, while the mean *G*_*s*_ is 9.5 *μ*N/m. There is a somewhat larger variation about the mean “_#_ of the normal RBCs in our experiments, which points to the heterogeneity of the population. It should be noted that our goal is not just to measure the mean *G*_*s*_ but also to understand how the values are distributed about the mean. Micropipette aspiration experiments report the value of the shear modulus to be 6 **–** 9 *μ*N/m^30^. Henon and others report a somewhat lower shear modulus value of 2.5 ± 0.4 *μ*N/m for healthy RBCS from their optical tweezer experiments^29^. Magnetic twisting cytometry^31^ measures the shear modulus to be between 6 and 12 *μ*N/m, while membrane fluctuation measurements^3^ report a value of **∼** 7 *μ*N/m.

**Figure 3.**
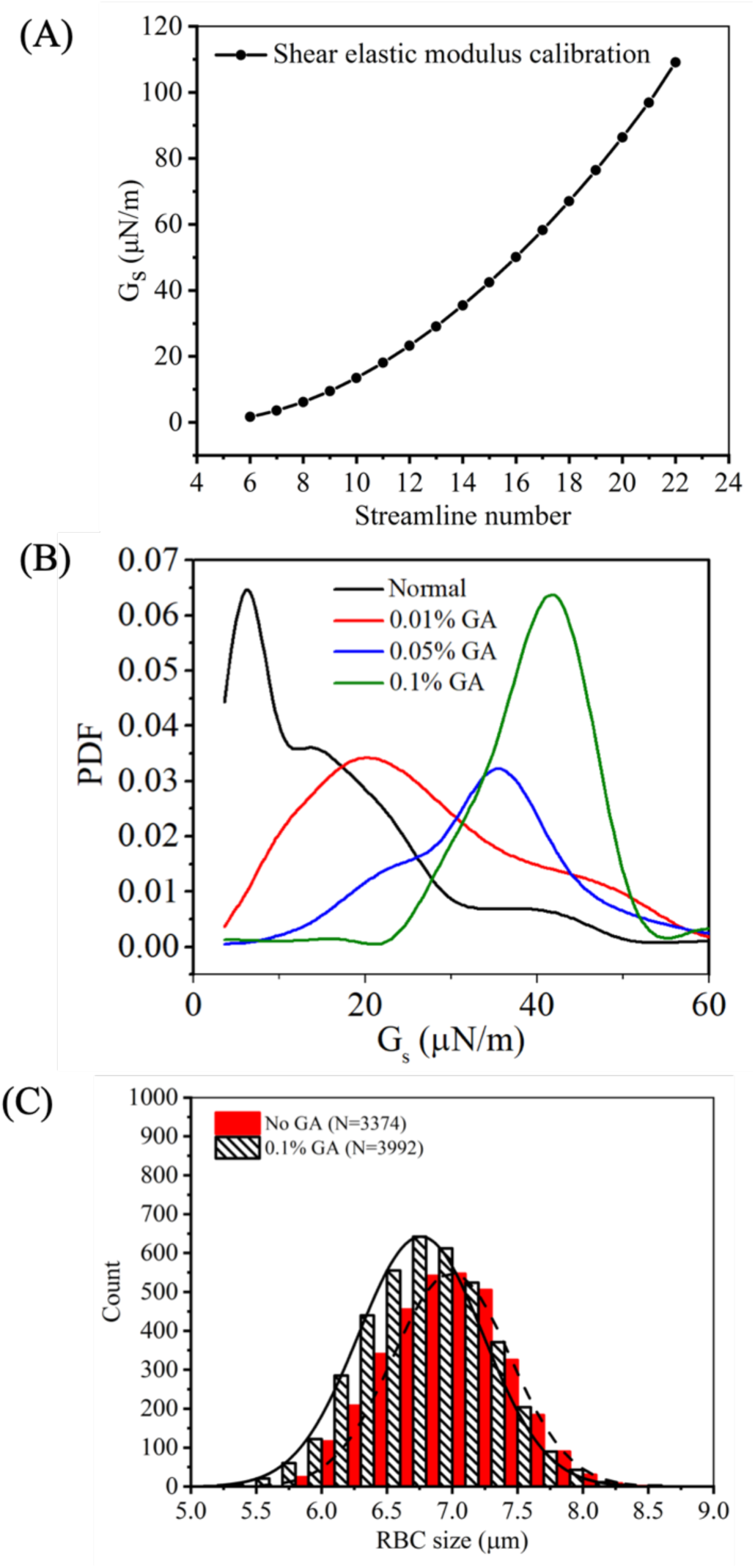
Shear modulus (G_s_) of normal and artificially stiffened populations of RBCs. (A) Calibration curve mapping each Gs to a specific streamline. G_s_ is calculated from equation (1) using the pressure values obtained from the simulation. (B) Probability density function (PDF) of G_s_ for normal and stiff populations of RBCs. The RBCs are stiffened by treating them with different concentrations of glutaraldehyde (GA) varying from 0.01% to 0.1%. The PDFs are normalized by equating the area under each curve to unity. (C) Size distribution of normal RBCs and the RBCs treated with the highest concentration of GA. N indicates the number of RBCs measured under each experimental condition. The size distribution does not change significantly with GA treatment.

We next analyze the data from experiments with chemically stiffened RBCs to generate the PDFs of *G*_*s*_ for these populations (fig. 3B). We find that the peaks of the PDFs shift towards progressively higher values with increase in GA concentration. The peaks occur at 15.8 *μ*N/m, *μ*N/m, and 38.9 *μ*N/m respectively. The mean values are 9.5 *μ*N/m (normal), 18.6 *μ*N/m (0.01% GA), 26.4 *μ*N/m (0.05% GA), and 32.1 *μ*N/m (0.1% GA) respectively. As expected, the mean *G*_*s*_ increases with an increase in the GA concentration. Furthermore, one or more knees appear in some of the PDFs. We speculate that this could be due to the non-linear elastic properties appearing at large deformations, perhaps showing the non-linear response of the spectrin network. The non-linear response (i.e. the values of *G*_*s*_ at which these knees appear) is slightly different for RBCs stiffened to different degrees. While GA is widely used in biology to fix and stiffen RBCs ^9,24^, at the moment there are no reported systematic characterization of the elastic constants of RBCs after treatment with specific concentrations of GA. Our results provide a way to controllably characterize the stiffness of RBC populations after GA treatment.

Note that the method we have used to obtain the calibration curve has a fixed parameter, the diameter of the biconcave undeformed RBC. If this changes significantly due to treatment by GA we must change the calibration curve accordingly. We measured the size distributions of normal RBCs (N = 3374) and RBCs stiffened with 0.1% of glutaraldehyde (N = 3992). We find that the size distribution of the RBC population does not change significantly even after treating them with the highest concentration of GA (0.1 %) used in these experiments (fig. 3C).

#### 2.2.2 Measurement of the Young’s modulus

We next measure the Young’s modulus of the RBCs. If *P*_1_ and *P*_2_ are the fluid pressures on the two curved faces of the annular sector representing the deformed RBC in our second simulation, then the average pressure on the RBC is given by 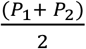. The undeformed diameter *D* of the RBC is assumed to be 7*μ*m in our calculations. Since the deformed RBC has a width given by *2r*_2_ the change in the linear dimension of the RBC under deformation is (*D* - *2r*_2_) Therefore, the Young’s modulus of the RBC (*Y*) is given by.

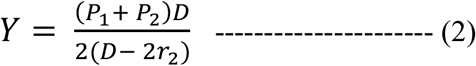

We follow the steps described in the previous section to generate a calibration curve (fig. 4A) of Young’s modulus vs. the streamline number. Combining our experiments with simulations, we can generate PDFs similar to the case of the shear modulus. The peak of the PDF lies at 5 kPa for normal RBCs. There are several reports on the measurement of Young’s modulus of healthy RBCs using AFM. On one hand Dulinska and others^9^ report values of Y varying from 0 to 40 kPa, with the mean value as 26 ± 7 kPa. On the other hand, Barns et al^24^ find the Young’s modulus to be 7.57 ± 3.25 kPa.

**Figure 4.**
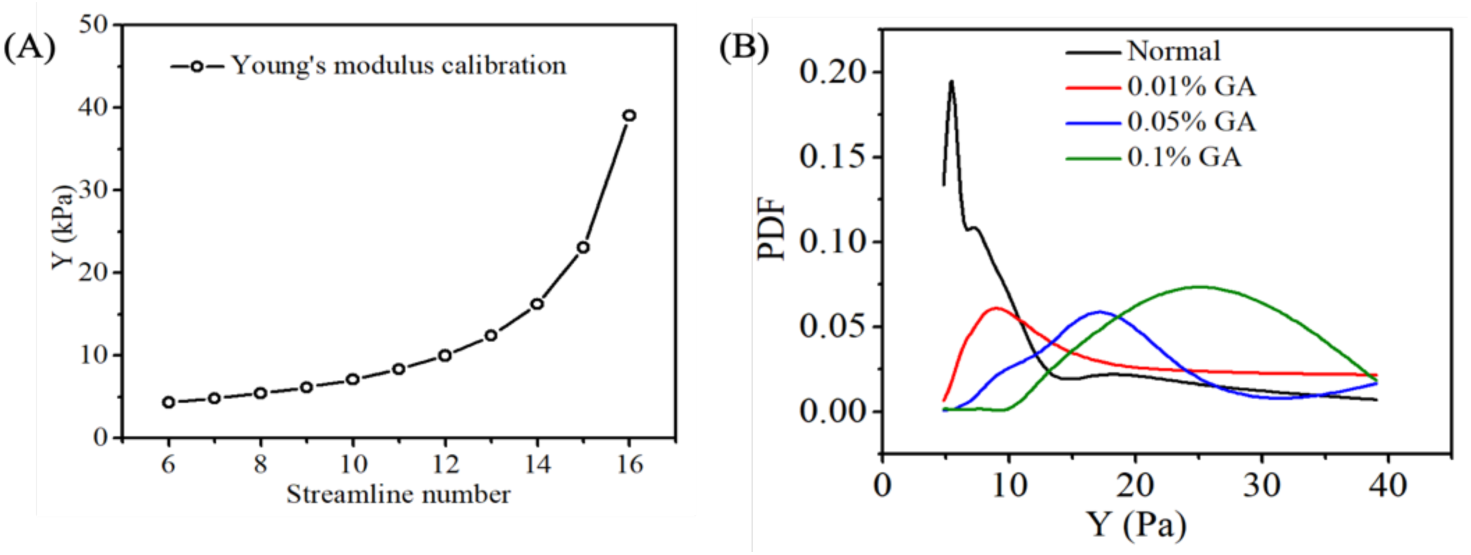
Young’s modulus (Y) of normal and artificially stiffened populations of RBCs. (A) Calibration curve to map each value of % uniquely to a specific streamline. (B) Probability density function (PDF) for the Young’s modulus of normal and stiff RBC populations. The peaks of the PDFs shift towards higher values for populations treated with increasing concentrations of GA. The PDFs are normalized by equating the area under each curve to unity.

We next obtain the distribution of *Y* for the three populations of RBCs treated with different concentrations of GA (fig. 4B). The peaks of the distributions shift to the right with increasing GA concentrations, and lie at 7.7 kPa, 14.3 kPa, and 19.6 kPa respectively. The mean *Y* for normal RBCs is 6.7 kPa, while the mean values for the increasingly stiffer populations are 10.2 kPa, 13.5 kPa, and 15.9 kPa respectively.

#### 2.2.3 Measurement of the Elongation Index of normal and stiff RBCs

The biconcave shape of mature RBCs has an important role in determining its deformability due to the reduced bending energy associated with this shape^32^. One of the most common indirect measures of the stiffness of a RBC is the change in its shape, given by the aspect ratio^23^ or the Elongation Index (*EI*) Both these quantities are measured by taking high resolution images of RBCs deformed by the same stress and then fitting an ellipse to the images. The *EI* of an ellipse with axes ‘*a*’ and ‘*b*’ is defined by:

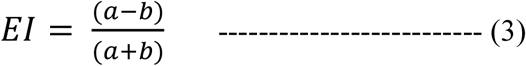

where ‘*a*’ and ‘*b*’ are respectively the semi-major and semi-minor axes of the ellipse. Clearly, it does not make any sense to compare the *EI* across different experiments. But it is useful as a rough estimate of the variation of deformability of RBCs within a given population, all of which are studied with exactly the same setup. The resolution and speed of our camera do not allow us to capture the exact shape change of an RBC as it goes past the obstacle. Nevertheless, we can provide an indirect measure of *EI* in the following manner. As we have explained before, with every streamline we associate a deformed shape of the RBC. This shape is not an ellipse but an annular sector as shown in fig.2B. We can now construct an ellipse with one axis equal to the width (*r*_2_) of this sector. The other axis of this ellipse is set by assuming that the area of the ellipse is equal to the area of the disk shape of the undeformed RBC. We can thus associate an *EI* with every streamline, and therefore, with every RBC that exits the device along that streamline. From this association, we can calculate the PDF of the *EI*s of both normal and hardened RBCs as shown in fig. 5. The *EI* is close to zero for ellipses which are close to circles in shape. Hence, we expect that the stiffer is a population of RBCs, the smaller the *EI* will be. This qualitative picture is confirmed in fig. 5 where we find that the peak of the *EI* for normal RBCs appears at **∼** 0.61. The peaks of the PDFs for RBC populations artificially stiffened by GA occur at progressively lower values of *EI* with increasing concentrations of GA. The peak appears at **∼** 0.2 for the highest concentration (0.1%) of GA. Similarly, the mean values of the *EI* occur at 0.61 (normal), 0.45 (0.01% GA), 0.34 (0.05% GA), and 0.26 (0.1% GA) respectively, thereby confirming the same trend.

**Fig. 5.**
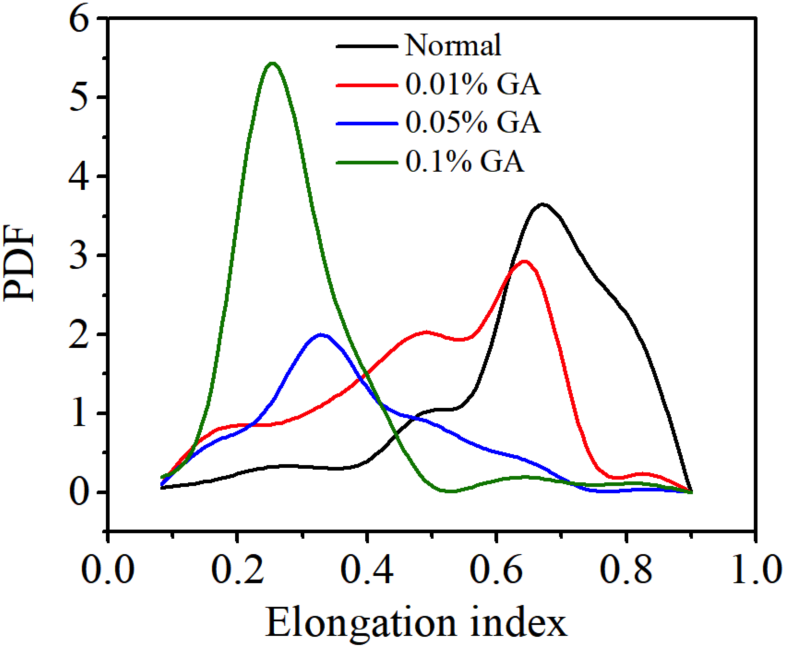
PDFs of the Elongation Indices (EI) of normal and hardened RBCs EI values are and higher for normal RBCs and progressively lower for stiffer populations. All PDFs are normalized.

### 2.3 Sorting of RBCs from a mixed population based on their deformability

Finally, we pass a mixture of 50% normal RBCs and 50% chemically stiffened (with 0.1% GA) RBCs through our device. We track their trajectories as they move through the device and calculate the PDFs for *G*_*s*_, *Y* and *EI* from the streamlines that they follow. As shown in fig. 6, all the PDFs are bimodal. Moreover, the peaks of all the PDFs for the mixed sample occur at values very close to the corresponding values we obtain for the individual populations. These results suggest a potential use of this device for sorting RBC populations based on their deformability alone.

**Fig. 6.**
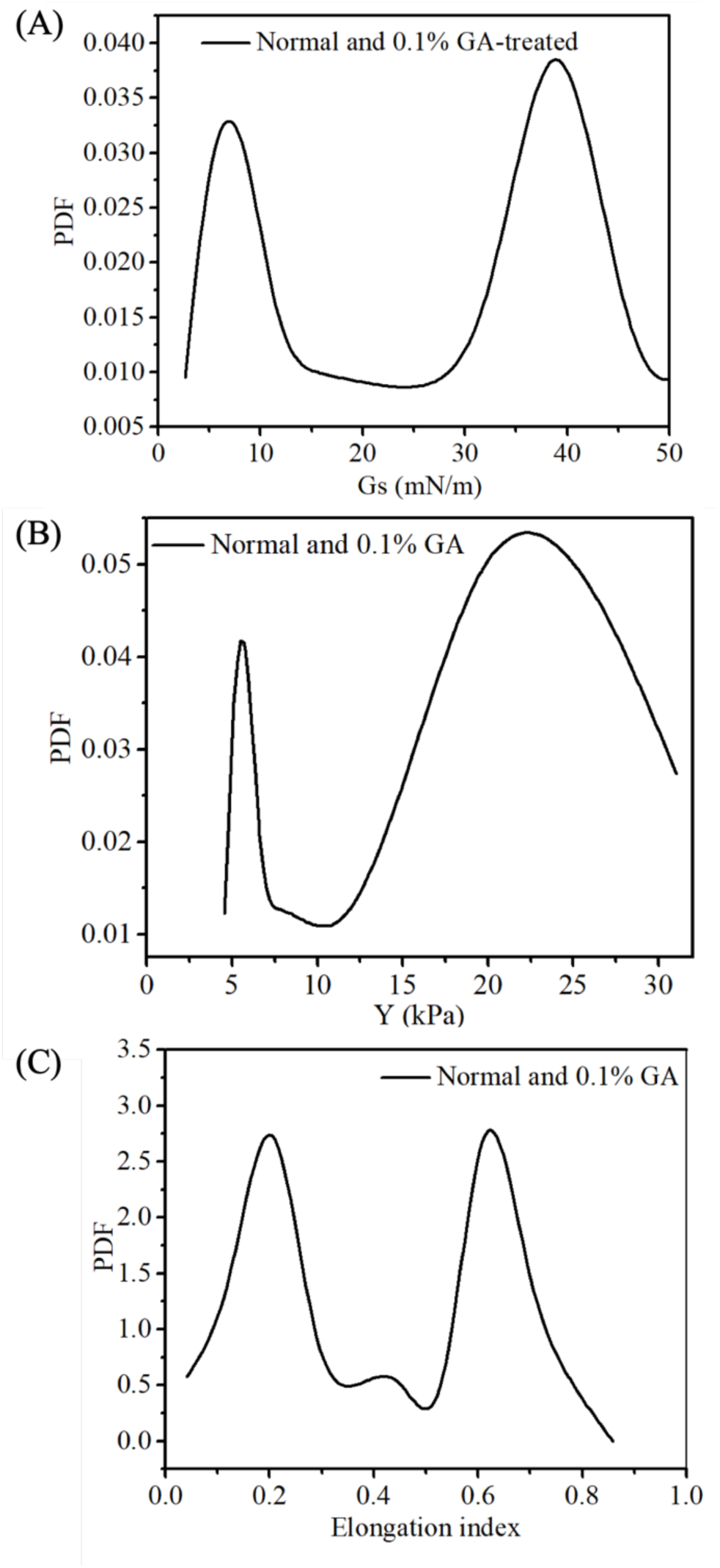
Experiments with a mixed population of RBCs obtained by mixing 50% normal and 50% chemically stiffened RBCs (0.1% GA). The distributions are obtained by analyzing 33058 cell tracks. (A) Bimodal distribution of G_s_ obtained from the mixed sample. (B) Bimodal distribution of % for the same sample. (C) Primarily bimodal distribution of EI.

## 3. Conclusions

We report a microfluidic device to obtain the distribution of the elastic constants (shear modulus, *G*_*s*_ and Young’s modulus, *Y*) of normal and stiffened RBC populations. We can also obtain the Elongation Index, *EI* of RBCs as a surrogate measure of deformability. The mean elastic moduli of normal RBCs obtained by us are in the same range as those obtained by other methods, such as, micropipette aspiration, optical tweezers, and AFM. We also pass a mixed sample containing equal numbers of normal and stiff RBCs through our device, and demonstrate that the PDFs of various elastic moduli have a clear bimodal distribution, with peaks at very similar locations compared to the ones obtained with individual populations. Thus, we can use our device to separate populations of RBCs based on deformability. Furthermore, our microfluidic device does not need fabrication of any feature size smaller than 10 *μ*m. It can work with a regular 25 fps microscope camera to measure **∼** 25,000 RBCs in each experiment at an average rate of 10 cells/s. Therefore, this method can be an inexpensive and fast alternative to the other deformability cytometry technologies currently available for measurement of elastic constants of red blood cells. It is ideally suited to construct a probability distribution function of deformabilities of a family of RBCs. Finally, our techniques can potentially be used to both measure and sort *extracellular vesicles* by size and deformability. These are about hundred times smaller than RBCs and lie at the limit of resolution of optical microscopes. They can be observed as bright point sources under high resolution microscopes, but their shapes under stress cannot be imaged by the existing techniques. So conventional deformability-based cytometry^14^ cannot be used to measure their elastic moduli. As our method relies measuring the trajectory of the elastic object, and not its shape, it is more easily generalizable to nanoscale objects.

## 4. Materials and methods

### 4.1. Design of the device

Our microfluidic device design (fig. 1A) is similar to that reported by Vesperini and others^17^. It has three functional sections for flow focusing, deformability measurement and collection of the sample. The first section focuses the stream of RBCs to the centre of the channel. The second section consists of a 40 μm-wide straight channel opening into a 45° funnel, with a single semi-circular pillar of 20 μm diameter placed at the mouth of the funnel. The flat side of the pillar faces the funnel, while the curved side faces the focused stream of RBCs. The third section has eight outlets, separated from each other by 18°, at the end of the funnel to collect the cell fractions following different streamlines. Our design has a few minor differences from that reported by Vesperini et al^17^. First, the gap between the obstacle and the channel wall is 10 μm in our device, which is larger than the diameter (∼7 μm) of a typical RBC. Second, the device height is 5 μm to ensure that the RBCs approach the pillar in a flat orientation instead of flipping sideways. Allowing for a larger height leads to flipping of the RBCs, which is avoided in our device design. Third, there are eight outlets instead of five for sample collection.

### 4.2 Microfluidic device fabrication

The microfluidic devices are made from the elastomer PDMS (Sylgard 184 from Dow Corning) by soft lithography^33^. We follow our reported protocol^34^ to make SU-8 masters for lithography with a height of 5 μm. We then mix the two parts of PDMS in 1:10 ratio, pour it on the master and cure in an oven at 65^°^C for 45 min. We punch inlets and outlets in the cured PDMS chip using a 1.5 mm diameter biopsy punch (obtained from Med Morphosis), and bond the chips to a 25 mm × 75 mm × 1.5 mm glass slide (Blue Star) using oxygen plasma (PDC 32G from Harrick Plasma) for 90 sec. The PDMS devices are used without any surface treatment.

### 4.3 Preparation of blood sample

We collect a drop of blood obtained by a finger prick from each healthy volunteer after obtaining informed consent. We dilute the blood by 1000X, either in normal saline (for untreated RBCs) or in glutaraldehyde (GA) solutions prepared in normal saline (for chemically stiffened RBCs). We treat blood with 0.01%, 0.05% and 0.1% concentrations of GA for 20 min at room temperature to stiffen them to different extents. To study the mixed samples in our device, we first prepare 2.5 ml of 0.1% GA-treated blood and mix them with 2.5 ml of untreated blood to obtain 5 ml of the mixed sample.

### 4.4 Experiment to measure the exit positions of the RBCs in the microfluidic chip

The blood sample is first passed through RAPID, a microfluidic cell sorting chip ^34^, to exclude WBCs and cell aggregates, and obtain a population of primarily RBCs. The RBCs collected from RAPID is introduced into our device for deformability measurement. Since the blood is diluted by 1000X, the presence of platelets does not affect our results. A focused stream of RBCs is allowed to flow towards the obstacle at a flow rate of 1 μl/min. After passing the obstacle, the RBCs leave the device through one of the eight outlets. To track the trajectory of the RBCs near the obstacle, we mount the microfluidic chips on a Nikon Eclipse Ti inverted microscope fitted with a 20X (0.45 NA) objective lens. The videos are acquired at a resolution of 640 × 512 pixels in uncompressed format by a 25 fps CCD camera, while keeping the obstacle at the centre of the frame. The details of the video processing algorithm in MATLAB to match the RBC tracks obtained from the experiments with the streamlines obtained from the COMSOL simulations are discussed in the Supplementary Information.

### 4.5 COMSOL simulation

We use the built-in microfluidics module (single-phase laminar flow) of the COMSOL multiphysics software (version 5.2) with ‘no slip’ boundary conditions and a fine mesh size. The simulations are performed with a flow rate of 1 μl/min to match the experimental conditions. Since the blood samples are diluted in saline by 1000 times, we simulate the flow of water to mimic blood flow in our device.

## Video processing to obtain cell tracks

We extract all frames of the video and use a MATLAB image processing algorithm to obtain the cell tracks. We read and store all frames as a 4D matrix. The number of the frames and their spatial resolution are then determined. We convert each frame from RGB to grayscale as we use a monochrome camera in our experiment and store the first frame of each video (fig. S1(A)) separately. This is done to calculate what alignment operations (e.g. rotation, scaling and translation as shown in fig. S1(B) and (C)) are needed on the image to overlap it with the streamlines obtained from the COMSOL simulation of the background flow. As indicated by the red dots on the inset of fig. S1(A), we use three coordinates on the obstacle to calculate the alignment factors--(*x*_1_,*y*_1_) and (*x*_2_,*y*_2_) are the coordinates of the left and the right corners of the obstacle respectively. Along with these two coordinates, we use the coordinate (*x*_3_,*y*_3_) of the center of the curved face to check the orientation of the obstacle.

Equations (1) and (2) show the formulae we use for image rotation, where (*x,y*) are the coordinates of the pixel to be rotated and (*xCos*Φ − ySinΦ, xSinΦ + ySinΦ) are the new coordinates of the same pixel after rotation.

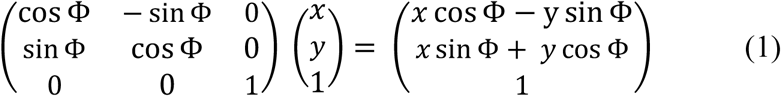

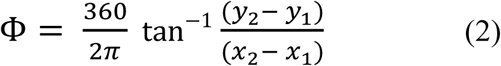

The rotation angle Φ is given by equation (2). If *y*_3_ < *y*_2_ a rotation angle (Φ + 180) is used. Equation (3) gives the scale factor needed to resize the image so that it fits on the COMSOL simulation window. The Scale Factor (SF) is the ratio of the distance (*x*_*c*2_ − *x*_*c*1_)between the two corners of the obstacle in the COMSOL simulation to the distance (*x*_2_ − *x*_1_)between the same two points in the image.

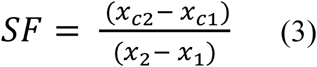

After rotating and scaling the image, we match the coordinates (*x*_*c*1,_*y*_*c*1_) in the COMSOL image (fig. S1(D)) to the corresponding point (*x*_1_,*y*_1_) in the video frame by a translation operation. Finally, we crop the image and overlap it with the COMSOL simulation window as shown in fig. S1(E).

To generate the RBC trajectory, we subtract two consecutive frames. This difference gives us the overlapping areas of a RBC at two different time points as shown in fig. S1(F). The difference image is then converted from greyscale to a binary image using a threshold value of 0.05. We then check whether the feature in the difference image (panel F) thus obtained is indeed a RBC by measuring its area. It is taken to be a RBC if its area is more than 50 pixels (corresponding to 6 μm diameter). The streak is then filled using morphological operations and its centroid is determined. The position of the centroid is next multiplied with all possible streamline patterns generated by COMSOL. The only streamline corresponding to a non-zero product after multiplication is chosen as the correct streamline indicating the track of the RBC.

We ignore any tracks mapped to streamlines 1 to 5 as the radius (*r*2) of the objects following these streamlines is too small (**∼**1 μm) to be RBCs. Similarly, all streamlines higher than 23 correspond to objects whose sizes are too large (**∼** 4.6 μm) to be RBCs. We then divide the entire range between streamlines 6 and 22 into several bins. The bin size is set by the resolution (0.9 μm) of our image acquisition system.

**Figure S1.**
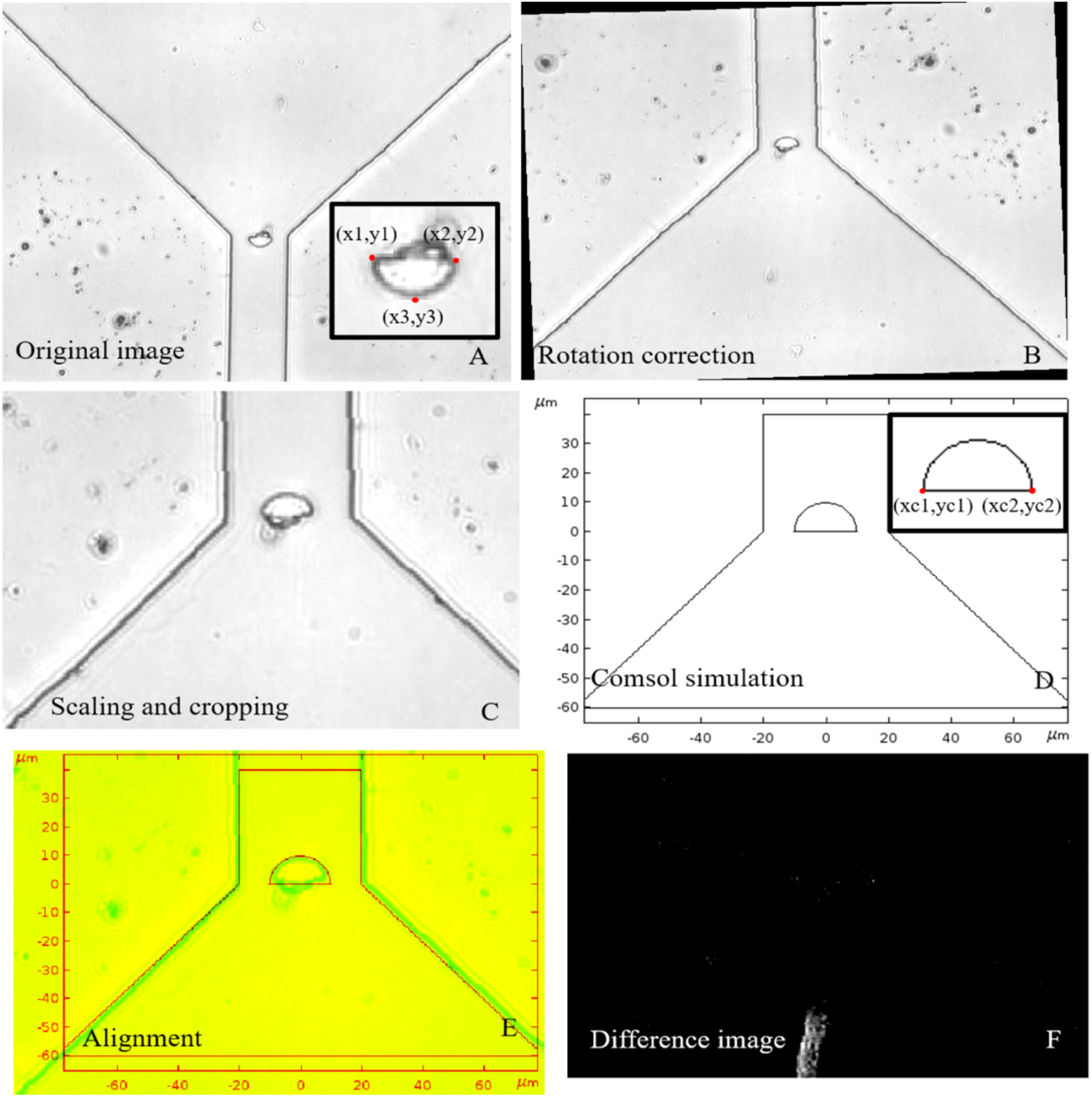
Video processing algorithm. (A) A frame from the original video acquired with a monochrome, 8-bit, 25 fps camera with a resolution of 640 × 512 pixels. The snippet shows a magnified view of the obstacle. Three points marked in red on the obstacle are used to correct the alignment of this image with the streamlines obtained from COMSOL simulation of the background flow. (B) The image is rotated to remove any angular displacement. (C) Scaling and cropping operations resize each frame of the video to the size of the COMSOL simulation window shown in (D). (E) Each frame of the video is overlapped with the COMSOL simulation window to perform a binary multiplication. (F) The difference image is computed from the current and the previous frames to identify moving objects. The outline of a RBC can be seen as a streak in (F). This area of this outline is then checked for the correct size range of a RBC and the coordinates of its center of mass is obtained. Finally, matrix multiplication is used to determine the position of that particular center of mass on the corresponding streamline.

